# An Open-Source Pipeline for Analyzing Changes In Microglial Morphology

**DOI:** 10.1101/2021.01.12.426422

**Authors:** Devin Clarke, Hans S. Crombag, Catherine N. Hall

**Author notes:** Corresponding author’s address.

## Abstract

Changes in microglial morphology are powerful indicators of the inflammatory state of the brain. Here we provide an open-source microglia morphology analysis pipeline that first cleans and registers images of microglia, before extracting 62 parameters describing microglial morphology. It then compares control and “inflammation” training data and uses dimensionality reduction to generate a single metric of morphological change (an ‘inflammation index’). This index can then be calculated for test data to assess inflammation, as we demonstrate by investigating the effect of short-term high fat diet consumption in heterozygous Cx3CR1-GFP mice, finding no significant effects of diet. Our pipeline represents the first open-source microglia morphology pipeline combining semi-automated image processing and dimensionality reduction. It uses free software (ImageJ and R) and can be applied to a wide variety of experimental paradigms. We anticipate it will enable others to more easily take advantage of the powerful insights microglial morphology analysis provides.

## INTRODUCTION

Microglia, the brain’s resident immune cells, are involved in phagocytosis and regulation of the adaptive immune response (Aloisi, 2001; Wolf et al., 2017). While resting they have a small soma and long, dynamic processes, allowing them to survey their local environment (Tremblay et al., 2011) for pathogens or damage associated molecular patterns (DAMPs; Colonna and Butovsky, 2017). Such stimuli activate a morphological shift towards an amoeboid-like shape which facilitates migration to sites of injury and phagocytosis (Nimmerjahn et al., 2005; Tremblay et al., 2011). To study such changes in morphology *in vivo*, transgenic animals expressing endogenous fluorescent reporters in microglia can be imaged through a cranial window over time (Hierro-Bujalance et al., (2018) for a review of multiphoton *in vivo* imaging of microglia).

However, analyzing morphological changes is often done manually. This can lead to rater error, demands a large time investment, and limits the number of cells that can be analysed. Furthermore, selecting which metrics of microglial anatomy to measure, from the number of branches to the overall shape of the soma, can be highly arbitrary. Although measuring many parameters simultaneously might better capture subtle variations, the multiple comparisons involved increase the chance of a false positive, whilst correcting for these decreases statistical power. With complex studies demanding large sample sizes to detect small effects, a more efficient approach is necessary.

As such, the three desirable features of a microglial morphology analysis pipeline are: 1) automation to reduce rater error and time investment, 2) measurement of multiple morphological features and 3) reduction in dimensionality of these features into a single index of morphological change.

Previously Kozlowski and Weimer (2012) produced an automated pipeline for extracting morphological data from *in vivo* microglial images, but only measured a limited subset of features and did not make their methods publicly available. Heindl et al. (2018) went a step further by combining a range of measurements into a single metric that tracked microglial activation. However, their image segmentation work was not appropriate for use with *in vivo* images, in which motion artefacts need to be corrected, and they did not provide access to their dimensionality reduction process.

To address these limitations, we built an open-source microglial morphology analysis pipeline using both ImageJ and R (Fig. 1). We used our ImageJ plugin to process *in vivo* images of resting and bacterial lipopolysaccharide (LPS) activated microglia from mice. The plugin cleans images, segments microglia from background, and extracts 62 morphological features for each cell. Following this, we used our R package to combine the features best at discriminating between LPS conditions into a single ‘inflammation index’. Here we demonstrate the use of this inflammation index in assessing the impact of high fat diet (HFD) feeding on microglial morphology in CX3CR1-GFP^+/-^ mice and provide a link to the GitHub repository (https://github.com/BrainEnergyLab/Inflammation-Index) where users can download the two packages necessary to perform this whole analysis pipeline.

**Fig 1.**
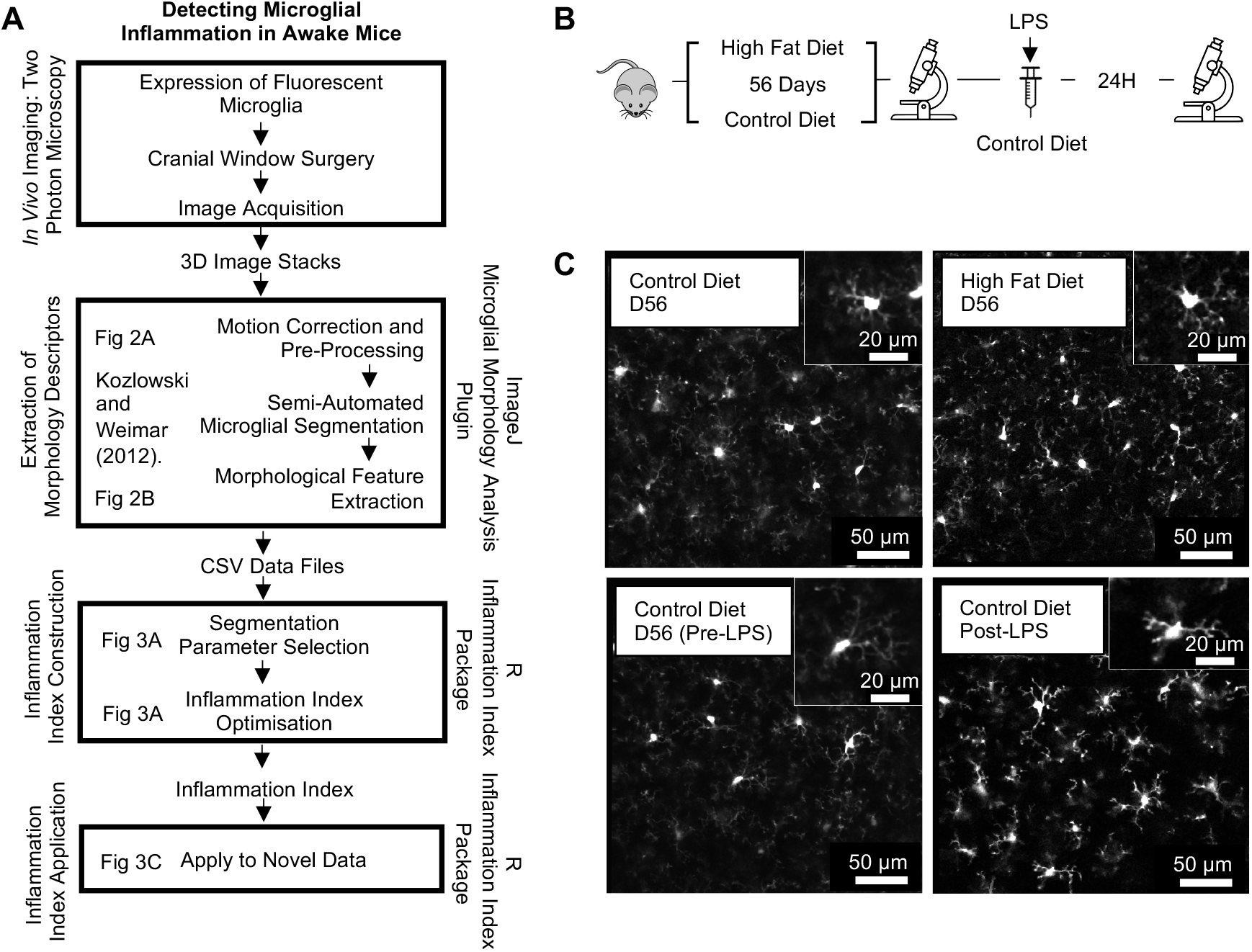
Experimental Design. (A) Inflammation index construction and application schematic. (B) Imaging timeline. Mice were imaged after 56 days of control or HFD feeding. Following this, control mice were injected with LPS and imaged 24 hours later. (C) Cx3CR1-GFP two-photon *in vivo* microglia images. Magnified view of example cells inset. Bottom row: pre- and post-LPS microglia (training dataset). Top row: microglia after 56 days of control or HFD (test dataset). Main scale bar represents 50 µm. Inset represents 10 µm.

## MATERIALS AND METHODS

First, we describe the experimental protocols used to generate our *in vivo* microglia datasets, before describing the detailed analysis pipeline, employing the ImageJ plugin and R package, that allows the level of microglial inflammation to be measured based on changes in their morphology (Fig. 1).

### Data Acquisition

#### Animals

Experiments were carried out in accordance with the guidelines of the UK Animals (Scientific Procedures) Act 1986. Mice (*Mus musculus Linnaeus*) were housed individually in a temperature-controlled room (22 ± 2°C) with a 12-hour light/dark cycle. A group of 10 mice, 12-18 weeks of age and of both sexes, were used with 5 mice fed a control chow diet (Research Diets D12450H-150FBZ, New Brunswick, NJ, USA) and the other 5 fed a HFD (nutrient balanced with 45% of total calories from fat, Research Diets D12451-150FBZ). Body weight was monitored daily. This sample size was chosen arbitrarily given we had no prior data based on inflammation index analyses to suppose an effect size. For diet group allocation, the first mouse that was available for imaging after cranial window surgery was allocated to the control group, and thereafter we alternated diet allocation for each next-available mouse. Data collected from these 10 mice was used in the test dataset, whilst data collected from the 5 control mice in pre- and post-LPS conditions formed the training dataset.

Mice were C57/BL6 and heterozygotes for Cx3CR1-GFP, so microglia expressed GFP (Jung et al., 2000). Heterozygotic mice were used as Cx3CR1-GFP homozygous mice are known to show a detrimental phenotype being resistant to the beneficial effects of environmental enrichment on neuronal plasticity, CA1 and hippocampal long-term potentiation, and Morris Water Maze performance (Maggi et al., 2011).

#### Surgical Window Implantation Procedure

Cranial windows were implanted to enable two-photon imaging of microglia in primary visual cortex (V1). Mice were anaesthetized in an induction chamber with 4% isoflurane (IsoFlo, Zoetis UK Limited, Leatherhead, Surrey, UK) until their breathing rate reached 1.5 Hz at which time the isoflurane concentration was decreased to 2% and the induction chamber was flushed for 4 seconds. The animals were then secured in a stereotactic frame (Kopf Instruments, Tuunga, CA, USA). Reflexes were checked to ensure adequate anaesthesia levels before administering sub-cutaneous injections of 0.9% saline (400 µL), the opioid analgesic buprenorphine (1.2 µg, 0.3 mg ml^-1^, Vetergesic, Ceva Animal Health, Amersham, Buckinghamshire, UK), the anti-inflammatory steroid dexamethasone (120 µg, 2 mg ml^-1^, Dexadreson, MSD Animal Health, Milton Keynes, Buckinghamshire, UK), and the non-steroidal anti-inflammatory meloxicam (6.2 µg, 0.5 mg ml^-1^, Metacam, Boehringer Ingelheim Animal Health, Bracknell, Birkshire, UK) to reduce dehydration, post-operative pain, and inflammation, respectively. Temperature was maintained at 37°C throughout using a homeothermic blanket (PhysioSuite, Kent Scientific Corporation, Torrington, CT, USA).

Hair over the skull was first trimmed using scissors, before the remaining hair was removed with hair removal cream (Veet, Reckitt Benckiser, Slough, Berkshire, UK). The exposed skin was cleaned with saline, then ethanol, and finally iodopovidone (Betadine, Mundipharma, Cambridge, Cambridgeshire, UK). The skin and periosteum over the skull were cut away and removed with small spring scissors and forceps (Fine Science Tools, Heidelberg, Germany) across the entire dorsal skull surface. During this process, any bleeding was stemmed using absorption spears (Fine Science Tools). The edges of the skin were then sealed to the skull with surgical cyanoacrylate (Vetbond, 3M, Bracknell, Berkshire, UK) before surgical calipers and a pen were used to mark the location of the craniotomy, as a circle with a 3 mm diameter overlaying the visual cortex (centered on a point 3.10 mm lateral to lambda, and 1.64 mm anterior of the lambdoid suture). The skull, excluding the marked region, was then roughened using a scalpel to create overlapping scores in perpendicular directions to aid cement and head plate adhesion, before being covered in surgical cyanoacrylate. The mouse was then tilted on the head mount so that the area marked for the craniotomy lay flat. The roughened area was covered with dental cement (Unifast TRAD, GC Europe, Leuven, Belgium; previously mixed with black acrylic) and a custom-built stainless-steel head plate (University of Sussex workshop, Brighton, East Sussex, United Kingdom) was placed over the dental cement and left for a few minutes until dry.

Following this, a dental drill (OmniDrill 35, World Precision Instruments, Sarasota, FL, USA) was used to drill around the edges of the marked area alternating between 0.5, 0.7, and 1 mm drill bits as necessary (diameter at the tip, Fine Science Tools) ensuring that regular breaks were taken and that the area was frequently irrigated with saline to prevent overheating of the brain (temperature increases at the brain’s surface of 25°C can result from continuous drilling, and are associated with significantly increased blood-brain barrier permeability. This did not occur when regular breaks were taken; Shoffstall et al., 2018). The area surrounding the marked area was also flattened. Once the skull was thin enough, the bone was moistened with saline for a final time and then lifted off with forceps and a microprobe. Gelfoam (Pfizer, New York, NY, USA) was used to stop any bleeding of the dura. Once bleeding was halted, forceps were used to remove the dura (if still present after the craniotomy).

An optical window (made from two 3 mm glass coverslips and a 5 mm glass coverslip (Harvard Apparatus, Holliston, MA, USA) sealed with optical adhesive (Norland, Cranbury, NJ, USA)) was placed into the craniotomy and secured using a glass rod whilst absorption spears were used to dry the liquid surrounding the window. The edges of the glass window were then sealed to the skull, first with surgical cyanoacrylate, then with dental cement. Finally, two rubber rings were secured on top of the head plate with dental cement to serve as a water reservoir during two-photon imaging using a water-based objective. Anaesthesia was removed and mice were placed into an incubator (37°C) to waken, before being singly housed in a recovery cage. For three days following the surgery mice were given daily 10 µg doses of meloxicam mixed into wet food mash.

#### *In vivo* Experimental Set-up

Beginning at least two weeks after surgery mice were gradually habituated to the imaging rig. Habituation reduces both excessive movement during image collection and chronic restraint stress, where the latter can influence microglial morphology (Hinwood etal., 2013). *In vivo* images were collected 12 weeks after surgery, well after astrocytic and microglial responses to the surgery had subsided (30 days after V1 cranial window surgery for astrocytes, and after 7-10 days for microglia; Holtmaat et al., 2009). During imaging sessions, mice were head-fixed underneath a two-photon microscope (Scientifica, Uckfield, UK) atop of a free-moving polystyrene cylinder (Biosciences Workshop, UCL) fitted with a rotary encoder (Kubler Group, Villingen-Schwenningen, Germany) that allowed recording of voluntary running. Imaging sessions were performed in the dark. Immediately prior to imaging sessions the vascular lumen was labelled either by intravenous injection with 2.5% (w/v) Texas Red Dextran (70 kDa, Invitrogen, Carlsbad, CA, USA) or subcutaneous injection of 2.5% (w/v) Texas Red Dextran (3 kDa, neutral, Invitrogen).

#### Two-Photon Microscopy

High-resolution imaging of microglia was performed with a commercial two-photon microscope (SP5, Leica Microsystems, Wetzlar, Germany) using a high numerical aperture water-immersion objective (Olympus 20X XLUMPLFLN20XW, Shinjuku City, Tokyo, Japan) with a working distance of 2mm, and a mode-locked Ti-sapphire infrared laser (Chameleon, Coherent, Santa Clara, CA, USA). Tissue was excited at a 940 nm wavelength, and the emitted light was filtered to collect green light from GFP (Cx3CR1-GFP labelled microglia). Imaging sessions were recorded using SciScan software (Scientifica). Laser power at the objective was kept below 25 mW to minimize photodamage. Laser power and photodetector gain was kept relatively constant between imaging sessions to minimise variations in the signal-to-noise ratio of images. Stacks sampling 297 x 297 x 100 µm of tissue (x, y, and z dimensions respectively) were collected from 50 to 150 µm below the window at a resolution of 512 x 512 pixels with a voxel size of 580 x 580 x 1000 nm (XYZ). Six frames were taken at each imaging plane, and image stacks were saved as single channel stacks in .tiff format. Imaging parameter choices were based on Kozlowski and Weimer (2012). Our ImageJ and R packages should work with images captured using different parameters but have only been verified with images captured using these specifications.

#### Lipopolysaccharide-Induced Microglial Activation

Around 12 weeks after cranial window surgery, mice were given intraperitoneal injections of LPS (O111:B4; Sigma-Aldrich, St. Louis, MO, USA) at a dosage of 4 mg kg^-1^ in 100 µl of 0.9% saline to induce microglial activation and imaged 24 hours later. Following the imaging session mice were culled using a terminal dose of intraperitoneal sodium pentobarbital (Dolethal; Vétoquinol UK Ltd, Towcester, Northamptonshire, UK) at 120 mg kg^-1^ (diluted to 10% in saline).

#### Dietary Manipulation

Dietary manipulation began no sooner than 4 weeks after cranial window surgery. Mice were fed either control diet or HFD for 56 days. Microglial images from V1 were acquired on day 56 of dietary manipulation. Control diet mice were subsequently injected with LPS as described above.

#### Image Processing and Analysis

Images were processed and analyzed using the Fiji distribution of ImageJ (Schindelin et al., 2012). By having a third party rename animal identifiers in the file and folder names all image processing was done blind to mouse identity and dietary or LPS treatment. The custom-written ImageJ plugin (Microglial Morphology Analysis; https://github.com/BrainEnergyLab/Inflammation-Index) performs all image processing steps as described below, which are automated except where indicated. Details of how to use the plugin, including the file structure required, can be found in the README file (https://github.com/BrainEnergyLab/Inflammation-Index/blob/master/Using%20the%20Microglia%20Morphology%20Analysis%20ImageJ%20Plugin.md).

#### Image Preprocessing

First, two-photon images needed to be preprocessed to improve image quality and remove motion artefacts as mice were imaged when awake (Fig. 2A). The order of the frames in the input stacks were of the structure F1Z1, F2Z1, F3Z1, F1Z2 etc., where each F denotes a single frame captured at each Z depth.

**Fig 2.**
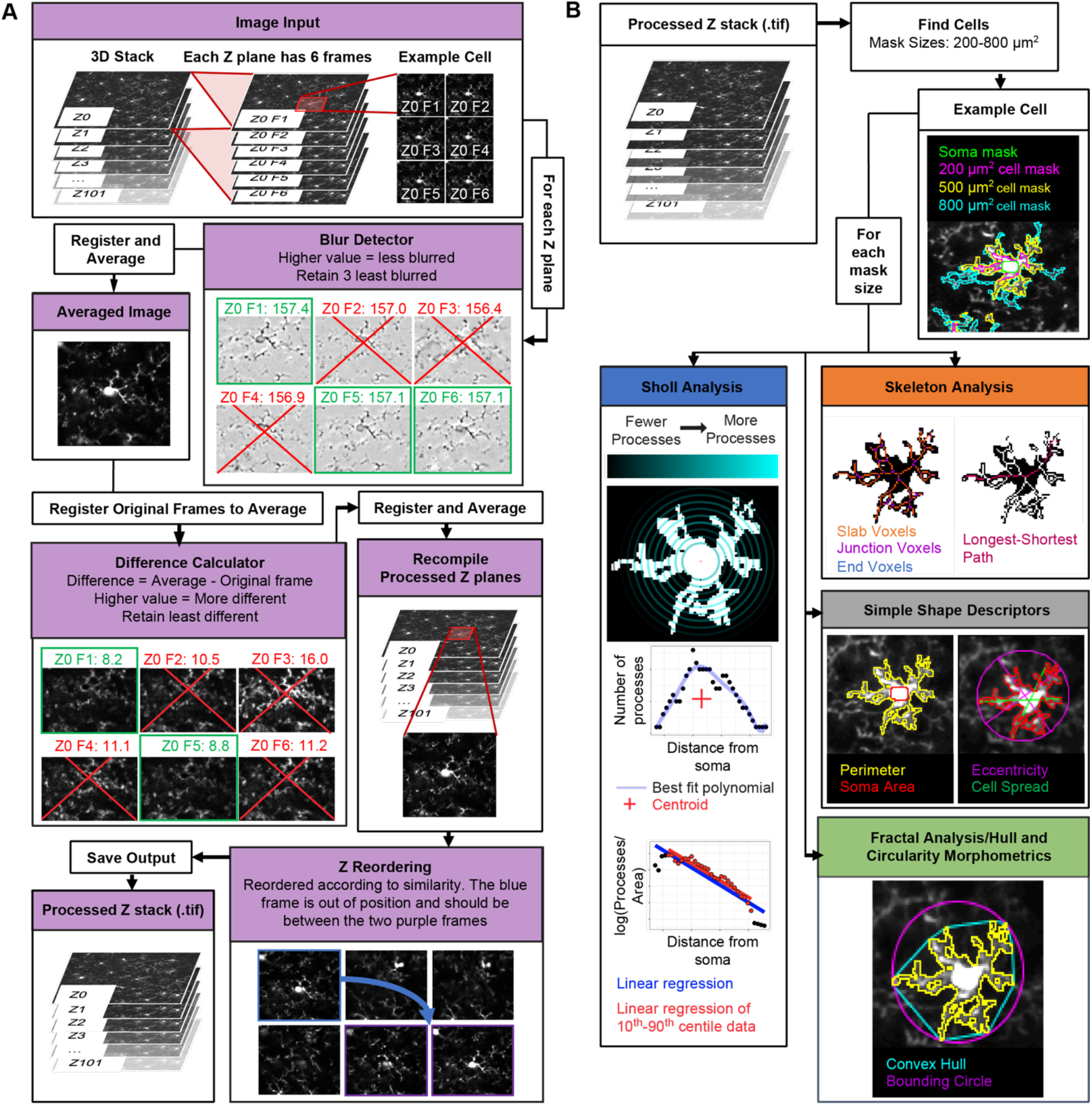
Image Processing and Extracting Morphological Descriptors. (A) Image processing schematic. Input stacks were split into substacks containing all frames from a distinct Z level. For each substack the least blurry frames were registered and averaged to create a reference frame that each frame of the substack was then registered to. The frames least different from the reference were again registered and averaged into a single frame. These single frames were stacked and reordered to represent their true positioning in Z. (B) Morphological feature extraction. After generating cell masks for a user-specified range of target mask sizes, the ImageJ plugin measured 62 different morphological features for each cell. These features were derived from five domains; simple shape descriptors, skeleton analyses, fractal analyses, hull and circularity morphometrics, and Sholl analyses.

a. To enhance the image clarity, contrast levels within a 3D stack were normalized using the Stack Contrast Adjustment plugin (Čapek et al., 2006) before the stack was divided into substacks of frames captured at each Z level (our image stacks represented 100 µm of tissue, and were split into 100 substacks, each containing 6 images).
b. Frames distorted by motion during image acquisition were then removed from each Z plane substack. Each frame within a substack was passed through a blur detector, where it was subject to a Laplacian of Gaussian (LoG) filter from the FeatureJ suite of ImageJ plugins (Meijering, 2002), and the maximum pixel intensity in the resulting image was measured. This filter highlights regions of an image containing rapid intensity changes (typically edges), with the maximum intensity in the resulting image an indicator of sharpness (where blurred images tend to have fewer edges). Using this method, the three least blurry frames in each substack were identified (users can set this number as they wish to fine tune their preprocessing steps). These were then registered to each other using the translation method in the MultiStackReg ImageJ plugin (Thévenaz et al., 1998) and averaged by creating a mean intensity projection of the three frames to yield a minimally blurry single image for each Z plane that served as a reference frame for detecting motion between frames within the Z plane substack. Frames with significant jitter were detected by registering each frame within the original substack to the reference frame (using the translation method in MultiStackReg) and calculating the mean difference between pixels in the reference image and each frame of the newly registered original substack using the Image Calculator functionality in ImageJ. The two least different frames were retained for each Z substack and were registered to each other (using the translation method in MultiStackReg; users can set this number as they wish). They were then averaged by creating a mean intensity projection to generate a cleaned frame for each Z plane. These cleaned images were recompiled to create a preprocessed and motion cleaned version of the input 3D stack, now composed of single frames representing each Z plane. The general approach applied in this step, i.e., selecting the least different frames relative to a reference frame for different Z substacks, was inspired by the work of Soulet et al. (2013).
c. Because movement in Z during image acquisition can mean that the actual Z location of acquired frames can differ from their apparent location, we next reordered each Z frame in the recompiled stack according to their actual Z location using the Z-spacing correction plugin (Hanslovsky etal., 2015).
d. Stacks were then inspected manually to detect failures in preprocessing (e.g., where Z reordering in step c) had failed, or where significant movement in the 3D stack was still present). In these cases, the least blurry and motion-distorted frames at each Z plane were selected manually from the original input 3D stack before being recompiled and registered. Where this manual selection was still not adequate (e.g., in cases where there was such severe movement during image acquisition that combining the highest quality frames for each Z plane still yielded too much movement or blurriness in the final stack), images were excluded from further analysis. These manual ‘quality control’ and frame selection steps are streamlined in the ImageJ plugin to make them as easy as possible for users.

#### Semi-Automated Microglial Segmentation and Quantification

The methodology used to semi-automatically generate microglial cell masks from two-photon image stacks is based on the work of Kozlowski and Weimer (2012). As with the image preprocessing steps, all the following steps were run using the custom-written ImageJ plugin: Microglial Morphology Analysis. In essence these steps build on the methodology described by Kozlowski and Weimar (2012) but extract a much greater array of morphology measurements from segmented cells. This process is built into the automated pipeline, with any manual input needed indicated clearly in the following steps.

a. Identification of cells: our 100 µm deep z-stacks were split into three 10 µm substacks separated by 20 µm (preventing the same cells being sampled in multiple substacks). Frames within each substack were averaged using a mean intensity projection. Cell locations were detected automatically (by identifying maxima in these projections using the Find Maxima plugin in ImageJ) before being either manually approved, or manually edited and approved, on these projections. Following this, a 120 x 120 µm region of interest (ROI) was drawn around each marked cell. The ROIs were thresholded with the ImageJ thresholding plugin using Otsu’s method (Otsu, 1979) to find the initial thresholding value. All pixels above this value that were in contact with the marked cell locations within each ROI were then used to form a cell mask by implementing the Find Connected Regions plug-in in ImageJ (Longair, 2006). The area of this mask was calculated and then compared to a user defined target area (from here on referred to as mask size). If the measured area of the mask matched the user defined mask size (within user defined limits - we used ± 100 µm^2^), the mask was considered complete. Otherwise, iterative thresholding was used whereby the applied threshold value was adjusted until either the measured mask area fell within the desired range, or the mask area stabilized for three consecutive iterations. If after passing these criteria a mask was within 5 µm of the edge of the ROI, it was rejected. The formula used to determine the threshold for the next iteration (Eqn 1) relates the threshold for the next iteration (*T*_1+1_) to the current threshold (*T*_1_), the area of the current threshold (*A*_1_), the mask size (*MS*), and the number of iterations thus far (*n*). Cell masks were excluded from further analysis if anaccepted mask contacted the edge of the 120 x 120 µm ROI.

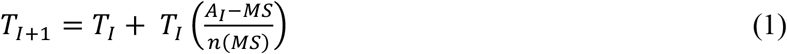
b. Soma detection: For each accepted mask, the cell soma were identified by first thresholding the ROIs using Otsu’s method (Otsu, 1979), then excluding particles that were smaller than 20 µm^2^ with a circularity value of less than 0.6. If a single particle remained, this was identified as the cell soma. These automatically generated soma masks were then subjected to user assessment, where if the mask was rejected, or automatic detection failed to identify a soma mask, a manual soma mask was drawn.
c. Measurement of morphological characteristics: Multiple parameters were calculated to capture different aspects of microglial morphology using a variety of Image J plugins (Fig. 2B). The Sholl Analysis plugin was used to perform Sholl analyses (Ferreira et al., 2014), the built in ImageJ measurement functions were used to extract simple shape descriptors (for example perimeter length, circularity), the built-in Skeletonize plugin and the AnalyzeSkeleton plugin (Arganda-Carreras et al., 2010) were used to analyse the skeletonized cell mask by, for example, calculating branch lengths, area occupied by the skeletonized cell, and the number of junction voxels in the skeleton, and FracLac (Karperien, 2013) was used to extract fractal, and hull and circularity morphometrics. The full list of extracted features is given in Table S1.

#### Composite Morphology Measure Construction

To best compare inflammation in different conditions, we combined all these measures into a single inflammation index, using an approach based on work by Heindl et al. (2018). This method identifies the morphological measures which best discriminate between conditions and passes these measures through a principal component analysis (PCA) to generate a single index of morphological change. In our pipeline, this process takes place using training data (in our case, pre- and post-LPS measurements), and the weights applied to the best discriminators to generate the single index are then applied to the test data (in our case control and HFD measurements) to evaluate if test conditions lead to significant changes in this morphological index (in our case, an index that reflects how similar to LPS-activated microglia a cell’s morphology is). However, as the data extracted from our images depend on the user-defined mask size, the results of this process will depend on what size is selected. Because of this, we first ran our image analysis steps on our pre- and post-LPS images for a range of mask sizes before constructing the composite index for each mask size. We then selected the optimal mask size for discriminating pre- and post-LPS cells and based our inflammation index on the index constructed using this mask size. We then extracted data from our test dataset (control and HFD images) using this mask size and applied the inflammation index to this. For this reason, inputting a greater range of mask sizes on which to run the initial microglial segmentation step maximizes the likelihood of creating a final inflammation index that is optimally sensitive to the training data conditions (in our case, pre- and post-LPS). Our R package ‘Inflammation-Index’ provides simple functions that users can employ to replicate this process, and like our ImageJ plugin is available at https://github.com/BrainEnergyLab/Inflammation-Index. The README for this package, including example inputs and outputs, can be found at https://github.com/BrainEnergyLab/Inflammation-Index/blob/master/Using%20the%20R%20Inflammation-Index%20Package.md. The R package uses the .csv results files output by the ImageJ plugin. As in the ImageJ plugin, the majority of steps are automated, with some input of user-specified arguments required. Only images from pre- and post-LPS-treated mice, previously fed a control diet, were analysed for construction of our inflammation index.

#### Mask Size Selection (Fig. 3A)

a. We ran our ImageJ plugin on our training data for a range of mask sizes (we used 200, 300, 400, 500, 600, 700, and 800 µm^2^ within limits of ± 100 µm^2^). For each mask size, receiver operating characteristic (ROC) analyses were conducted for each morphological measure to determine their ability to discriminate between activated and resting microglia (pre- and post-LPS treatment). ROC curves plot the false positive rate against the true positive rate of binary classification (i.e., pre- and post-LPS) at differing thresholds of a metric (i.e., for a morphological measure such as cell perimeter, we plot the false positive and true positive rates of classifying a cell as pre- or post-LPS at a given threshold of cell perimeter. We do this for all possible thresholds for our cell perimeter measurements). The area under the curve (AUC) of these plots tells us how well different measures discriminate between pre- and post-LPS microglia.
b. A selection of the most accurate discriminators based on their ROC-AUC values were retained. In this analysis we limited our selection to the best five to ensure we only included measures that were good discriminators, though users can set the number of measures to retain themselves and this number is optimized in later steps. The choice to use a set number of measures, rather than a set value of the AUC as a cutoff, is discussed later. A PCA with feature centering and scaling was run on these measures, with the first principal component (which by definition is the weighting of the inputs that best captures the variability in the data – in this case, variability due to inflammatory state) serving as an initial inflammation index for each mask size.
c. A number of our measures capture very similar features of a cell’s morphology. For example, some Sholl parameters are calculated multiple times for different subsets of the data (e.g., 10^th^– 90^th^ percentiles vs. the whole dataset), while hull and circle morphometrics are calculated based on both the centre of the bounding circle, and the centre of mass. In these cases, if multiple variants of the same metric were ranked as high discriminators, only the best performing variant was selected for inclusion in the PCA.
d. We then selected the mask size which most significantly discriminated between pre- and post-LPS conditions. The ‘Inflammation-Index’ R package allows two ways to identify the best mask size. The first, which we used for the analyses presented here, compares the p values of the difference in the inflammation index between training conditions (pre- and post-LPS) for each mask size (Fig. 3A). This comparison is done using a linear mixed model, with the animal identifier specified as a random intercept. The second option in the package calculates the AUC from a ROC plot of the inflammation index to select the mask size that offers the best discrimination between conditions (pre- and post-LPS). For our data the strongest effect of LPS was observed using a mask size of 400 µm^2^. This optimal mask size is detected automatically within the ‘Inflammation Index’ R Package, so all the following steps are performed only on data extracted using this mask size.

**Fig 3.**
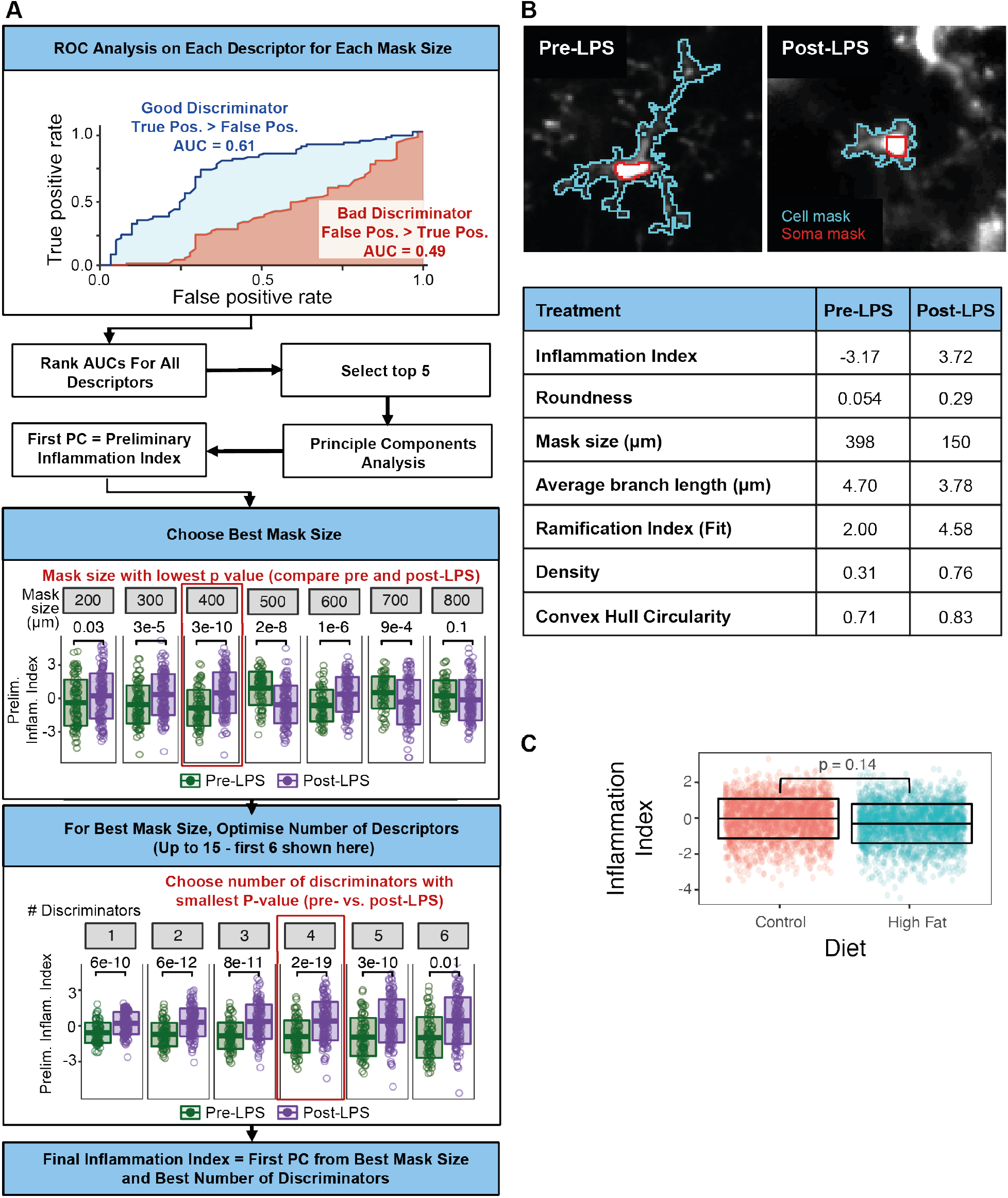
Calculating and Applying the Inflammation Index. (A) For each mask size, the morphological descriptors’ ability to discriminate between pre- and post-LPS conditions was evaluated using an ROC-AUC analysis (e.g. regression coefficient: blue, critical radius: red). A PCA was run on the top five discriminators, the first principal component compared pre- and post-LPS, and the mask size with the smallest p value selected. PCAs were then run on a range of the best discriminators at this mask size. The range (e.g. the best 4 discriminators) with the lowest p value for the effect of LPS was selected as the inflammation index. (B) Two example cells from the LPS dataset displaying values from the seven best discriminators. (C) This inflammation index can then be calculated for experimental data. It was unaffected by 56 days of HFD feeding. Dots are cells, lines represent the mean +/-1 s.d. 100 cells from 5 control mice, 138 cells from 5 HFD mice. Statistical tests were conducted using linear mixed models with animal ID specified as a random intercept to avoid pseudoreplication.

#### Inflammation Index Optimisation and Application

Following mask size selection, the next steps in the pipeline first refine the inflammation index to be optimally sensitive to the morphological changes associated with training conditions (whilst avoiding it being overly specific to the data it is based on). The optimized inflammation index can then be applied to novel data to test for any activation-associated microglial morphological change. Again, these steps are all automated though can be adjusted by user input where indicated.

a. Refining the inflammation index: The preliminary inflammation index is composed of a subjective number of the most accurate discriminating morphological measures (we used five). To optimize the inflammation index, we determined whether combining more, or fewer, measures produced a stronger effect of the training conditions (and therefore better discriminability between them). For the chosen mask size, the index was reconstructed using the best 1 to N discriminators of activated microglia using ROC curves as described above, where N is the user-defined maximum number of discriminators to be included. We used a maximum of 15 factors to avoid building an overly specific metric to the pre- and post-LPS data it was based on, and which therefore would be unfit for use with novel datasets. Inflammation indices derived using these different combinations of factors were then generated using a PCA as above and the best index was then identified as above by comparing p values of the effect of LPS treatment between indices (e.g., one constructed with three discriminators versus one constructed with four). The R package also offers users the ability to select this based on the AUC of ROC curves of each index.
b. This final inflammation index represents the first principal component of a PCA run on the optimal number of morphological discriminators. As such, its value is based on a weighted combination of the different discriminators that were included in its construction (for a comparison of two cells, illustrating the values of seven of the metrics identified as the best discriminators between pre- and post-LPS conditions, and the inflammation index, see Fig. 3B). As long as the same morphological measures are present, the same weighted combination can be computed for any novel dataset acquired in a similar way, and an inflammation index can be calculated. This inflammation index then allows users to evaluate the activation level of their microglia in different experimental conditions. The construction of this index for novel data is done automatically through the ‘Inflammation Index’ R package. The nature of the training data will affect what sort of changes will be best detected. As described here our inflammation index is optimized for detecting LPS-like microglial activation but use of other training data could allow optimal detection of, for example, morphological differences between peri-infarct or contralateral microglia in stroke models (as in Heindl et al., 2018).

## RESULTS AND DISCUSSION

We developed an open-source image analysis pipeline that facilitates the largely automated processing of large imaging datasets and the creation of a single metric of morphological change using dimensionality reduction to allow users to leverage *in vivo* microglial morphological analyses. This makes analysis easier, diminishes the effects of rater error on results, removes the need for assumptions about which metrics will be most sensitive to experimental manipulation, maximizes statistical power, and minimizes false discovery rates. Our ImageJ and R packages are freely available at: https://github.com/BrainEnergyLab/Inflammation-Index. A detailed description of their use is provided in the Methods section. Here we describe the construction of an inflammation index with this pipeline using data collected from mice before and 24h after LPS treatment (known to increase microglial activation). We then used this index to evaluate the inflammatory effects of 56 days of HFD feeding on microglial morphology (Thaler et al., 2012; Calvo-Ochoa et al., 2014; Denver et al., 2018).

### Image acquisition

Image stacks of GFP-labelled microglia were collected from V1 in awake CX3CR1-GFP^+/-^ mice using *in vivo* two-photon imaging (Fig. 1) after 56 days of either control diet or HFD consumption. After imaging, control mice were injected with LPS and imaged 24 hours later. The quality of these stacks depended on image resolution, frame capture rate, and motion during acquisition. We found a capture rate of around one frame per second with a resolution of 1.8 pixels per micron allowed us to acquire high quality images despite the mouse’s movement. Users of our plugins should trial different combinations of imaging parameters to produce the highest quality input given differences in imaging setups.

### Image Analysis

There are three broad stages to the analysis pipeline, which are described in full in the Methods section:

1. Cleaning of image stacks and extraction of morphology descriptors (Fig. 2)
2. Construction of a composite morphology index based on training data (our pre- and post-LPS images; Figs 3A and B)
3. Use of the composite index to assess morphological changes in test data (control and HFD images; Fig. 3C)

### Image Cleaning and Extraction of Morphology Descriptors

We used our ImageJ plugin to preprocess 3D image stacks and semi-automatically identify microglia and measure 62 morphological descriptors in our pre- and post-LPS dataset (Fig. 2).

#### Correction of Noise and Motion Contamination

For each Z location in a 3D image stack, our plugin uses an average image of the least blurry frames to identify the least motion contaminated frames, before averaging the latter to generate a single cleaned image. These single images are recompiled into a Z stack that is reordered to correct for shifts in Z position during image acquisition. Where noise or motion artefacts were too substantial to be corrected with these methods, image stacks were excluded from further analysis to only include the most accurate microglial renderings.

Our approach is based on the work of Soulet et al. (2013), who used an average projection of the frames in a Z plane as a template to detect, and remove, the frames in that plane that were most different from the template (i.e., contaminated by motion). We improved on this method by first adjusting the contrast across the entire stack, and then by using a blur detector to create our template using the least blurry frames. We implemented blur detection using the maximum grey value of images subjected to a LoG filter, adapting previous methods (Bansal et al., 2016) that used the variance of grey values, as we found the former was a better indicator of blurriness (Fig. S1).

#### Semi-Automated Microglia Segmentation & Morphological Feature Extraction

Following image cleaning we used our plugin to semi-automatically detect the location of microglia before generating cell masks using an iterative thresholding approach. Kozlowski and Weimar (2012) used iterative thresholding to segment microglia and demonstrated that masks generated using iterative thresholding had ∼90% similarity in terms of the numbers of processes counted when compared to manual results. Given that iterative thresholding requires a user-defined target mask size and limit, our R package can evaluate data collected using multiple mask sizes to detect the size most sensitive to morphological differences so that it can be used when processing test images. To take advantage of this, we computed masks using target sizes ranging from 200 to 800 µm^2^ inclusive at intervals of 100 µm^2^ (within a limit of ±100 µm^2^) for cells in our LPS dataset. To get as full a description of microglial morphology as possible, 62 morphological descriptors were derived from each cell mask, across five domains used previously to quantify microglial morphology (Kozlowski and Weimer, 2012; Fernández-Arjona et al., 2017). These were: shape descriptors (such as cell perimeter), skeleton analyses (for values such as the number of branches in a cell), Sholl analyses (for metrics based on the number of processes within a cell, such as the ramification index), hull and circularity morphometrics (such as the convex hull area) and fractal analysis (for values such as lacunarity).

### Construction of a Composite Morphology Metric

The data extracted by the ImageJ plugin was then used to build a single metric - the “Inflammation Index” - that was optimally sensitive to the morphological effects of LPS treatment with our R package (Figs 3A and B).

#### Mask Size Selection

Firstly, we used the R package to identify the mask size associated with the greatest sensitivity to morphological changes in the LPS dataset. This required computing an inflammation index for each mask size, which we based on the work of Heindl et al (2018). For each input mask size the morphological descriptors best at discriminating between the training data (i.e. LPS) conditions are identified using a ROC-AUC analysis. Whilst Heindl et al (2018) utilized a specific cutoff AUC value for selecting their best descriptors, we instead opted to use a cutoff based on their rank. As exact AUC values will differ depending on the quality of input images and the magnitude of the morphological changes associated with the training conditions, our approach allows greater generalizability across conditions, labs and experiments. A PCA is then run on these best descriptors with the first principal component from the PCA being used as the inflammation index. The index was then compared between training conditions and the mask size with the smallest p value for this comparison was identified as the optimal mask size. For our data this was 400 µm^2^ (Fig. 3A), though Kozlowski and Weimar (2012) found 500 µm^2^ (with a limit of ±100 µm^2^) provided maximal cell detection with good sensitivity to LPS-induced morphological changes. This is likely to differ across setups and experiments.

#### Inflammation Index Optimisation

Having defined the optimal mask size, the inflammation index was then refined into its final form by identifying the combination of morphological descriptors (i.e. the best one through four, one through five, one through six etc.) that led to the smallest p value between training conditions within the optimal mask size. For our data this was the best one through four features (roundness, mask size, average branch length, and the ramification index of the Sholl analysis linear regression). Our final inflammation index discriminated between our pre- and post-LPS training data with a standardized effect size of 0.654.

### Applying the Inflammation Index to Novel Data

The identity and weightings of the discriminators in the final index were then used to generate an inflammation index for microglia in our test dataset, where we imaged CX3CR1-GFP^+/-^ mice fed control diet or HFD after 56 days. These images were processed with the ImageJ plugin employing our optimal mask size (400 µm^2^ ±100) before the R package was used to apply the inflammation index generated using the LPS data to calculate an inflammation index for each test cell, effectively quantifying how similar the morphology of our test cells were to LPS-activated microglia.

Our results showed no effect of HFD feeding on microglial morphology (Fig. 3C). Though HFD has been shown to cause inflammation and activation of microglia in some strains of mice (Thaler et al., 2012; Calvo-Ochoa et al., 2014; Denver et al., 2018), the effects are expected to be weaker than those of LPS, which (unlike serum derived from mice fed a HFD for 16 weeks) significantly increases TNF alpha protein levels in isolated microglia (Baufeld et al., 2016). Cx3CR1-GFP^+/-^ mice possess a heterozygous knockout of Cx3CR1 function, which has recently been shown to make them resistant to inflammation, including after HFD feeding (Cope et al., 2018), so use of these mice likely resulted in us not observing HFD-mediated microglial activation. Of note, abuse of the inflammation index, by using control and HFD microglia as a training dataset to create an index that is optimally sensitive to the test data, produces a highly significant difference in a statistical test of control (mean ± s.d. of −1.004 ± 0.740) versus HFD (0.034 ± 1.113) conditions (p < 0.001; linear mixed model with animal ID specified as a random intercept). This highlights the importance of using independent test and training data sets.

### Conclusion

We present a fully open access analysis pipeline for analyzing changes in microglial morphology in large *in vivo* imaging datasets. Our two packages, in ImageJ and R, provide a streamlined process for analyzing complex morphological data and are available on GitHub to allow free access and encourage participation in their future development. The supplied packages can be applied to any experimental condition, e.g., detecting regional differences, simply by altering on the training data utilized. We anticipate our pipeline will greatly increase the ease through which multiple groups can take advantage of the power of *in vivo* microglial morphology analysis in a manner that is quick, streamlined, and preserves both statistical rigor and statistical power.

## ACKNOWLEDGEMENTS

We would like to thank K. Boyd for both setting up, and providing training on, cranial window surgery and two photon *in vivo* imaging in our lab. In addition, we’re grateful to L. Bell and the University of Sussex animal husbandry staff for maintaining our mouse colonies and looking after their welfare. A. Garnham provided insightful guidance on the statistical analyses to use, and the CAJAL Advanced Neuroscience Training Programme provided excellent training on *in vivo* microglial imaging and analysis. Finally, we’d like to acknowledge K. Shaw, O. Bonnar, and D. Grijseels, for their advice, assistance, and support throughout.

## AUTHOR CONTRIBUTIONS

D.C., H.S.C and C.N.H. conceived the study, D.C. collected data and designed and performed the analysis, under the supervision of C.N.H., D.C. and C.N.H. wrote the manuscript.

## LIST OF SYMBOLS AND ABBREVIATIONS USED

Cx3CR1: C-X3-C Motif Chemokine Receptor 1
GFP: Green Fluorescent Protein
DAMPs: Damage Associated Molecular Patterns
LPS: Lipopolysaccharide
HFD: High Fat Diet
V1: Primary Visual Cortex
LoG: Laplacian of Gaussian
ROI: Region of Interest
PCA: Principal Component Analysis
ROC: Receiver Operating Characteristic
AUC: Area Under the Curve

